# *In vivo* molecular signatures of cerebellar pathology in spinocerebellar ataxia type 3

**DOI:** 10.1101/2020.01.03.894337

**Authors:** Maria do Carmo Costa, Maria Radzwion, Hayley S. McLoughlin, Naila S. Ashraf, Svetlana Fischer, Vikram G. Shakkottai, Patrícia Maciel, Henry L. Paulson, Gülin Öz

## Abstract

**Background:** No treatment exists for the most common dominantly inherited ataxia Machado-Joseph disease, or spinocerebellar ataxia type 3 (SCA3). Successful evaluation of candidate therapeutics will be facilitated by validated noninvasive biomarkers of aspects of disease pathology recapitulated by animal models.

**Objective:** We sought to identify shared neurochemical signatures in two mouse models of SCA3 that reflect aspects of the human disease pathology.

**Methods:** Cerebellar neurochemical concentrations in homozygous YACMJD84.2 (Q84/Q84) and hemizygous CMVMJD135 (Q135) mice were measured by magnetic resonance spectroscopy at 9.4 tesla. Motivated by the shared neurochemical abnormalities in the two models, we determined the levels of neurofilament medium (NFL, indicator of neuroaxonal integrity) and myelin basic protein (MBP, indicator of myelination) in cerebellar lysates from a subset of mice and from patients with SCA3. Finally, NFL and MBP levels were measured in cerebellar extracts of Q84/Q84 mice upon sustained silencing of the mutant *ATXN3* gene from 6-8 weeks-of-age until death.

**Results:** Both Q84/Q84 and Q135 mice displayed lower *N*-acetylaspartate than wild-type littermates, indicating neuroaxonal loss/dysfunction, and lower *myo*-inositol and total choline, indicating disturbances in phospholipid membrane metabolism and demyelination. Cerebellar NFL and MBP levels were accordingly lower in both models as well as in the cerebellar cortex of patients with SCA3 than controls. Furthermore, long-term sustained RNAi-mediated reduction of ATXN3 levels increased NFL and MBP in Q84/Q84 cerebella.

**Conclusions:** *N*-acetylaspartate, *myo*-inositol and total choline levels in the cerebellum are candidate biomarkers of neuroaxonal and oligodendrocyte pathology in SCA3, which are reversible by reduction of mutant *ATXN3* levels.

## Introduction

With increasing human lifespan, numerous age-related neurodegenerative diseases of diverse etiologies are becoming more prevalent in our societies. Nine such disorders are caused by an expanded CAG triplet repeat in specific disease genes that encode an abnormally longer polyglutamine (polyQ) tract in the respective disease proteins and, therefore, are denoted as polyQ diseases^1, 2^. Since the identification of these disease genes in the 1990s, advances have led to increasing understanding of the molecular basis of polyQ disorders and numerous therapeutic agents have been tested in animal models. Due to the combined efforts of academy and industry, several potentially efficacious therapies have emerged for individuals with polyQ diseases. Nucleotide-based reduction of CAG-containing transcripts using nucleic acids has proven to be a particularly compelling therapeutic strategy for polyQ diseases in pre-clinical studies^3–7^, and clinical trials are underway for Huntington disease (HD) using *HD*-targeting antisense oligonucleotides (ASOs)^8^. While nucleic acid based therapy for polyQ disorders is very encouraging, other gene and pharmacological therapeutic approaches are also in the pipeline^9–11^. Readiness for these upcoming clinical trials depends, however, on the availability of disease biomarkers that allow noninvasive monitoring of aspects of cerebral disease pathology during the course of the trial to assess therapeutic efficacy.

Machado-Joseph disease (MJD) or spinocerebellar ataxia type 3 (SCA3) is a polyQ disease caused by an expanded CAG repeat in the *ATXN3* gene^12^ that leads to selective dysfunction and loss of neurons of specific nuclei in the cerebellum, brain stem, midbrain, spinal cord, and peripheral nerves^9–11, 13–21^. This central and peripheral nervous system pathology manifests in patients with SCA3 as ataxia and a variable degree of other symptoms, such as extrapyramidal signs, neuropathy, ophthalmoplegia, visual impairment and muscle atrophy^22–26^.

SCA3 is one of the polyQ diseases for which clinical trials are forthcoming. ASOs^27–30^, other RNA interference molecules^31, 32^, and pharmacological agents such as the selective serotonin reuptake inhibitor, citalopram^33–35^, showed preclinical efficacy to mitigate molecular, pathological, and phenotypic SCA3-like signs in transgenic mice.

The success of upcoming human trials of these agents will be facilitated by the availability of validated noninvasive biomarkers of cerebral and cerebellar pathology. To this end, a number of studies have demonstrated an ability to detect macro-and micro-structural and neurochemical abnormalities noninvasively by magnetic resonance imaging^14, 36–38^ (MRI) and spectroscopy^39–41^ (MRS) in patients with SCA3. While macrostructural disease-associated changes detectable by conventional MRI clearly mark tissue loss, other MR metrics may allow noninvasive monitoring of aspects of pathology that precede cell loss and atrophy, and hence may be reversible with treatments. However, such MR metrics need to be validated by comparing them to tissue pathology.

Here, using high sensitivity MRS methodology at 9.4 tesla (T), we sought to identify cerebellar neurochemical signatures in two mouse models of SCA3 that are extensively used in SCA3 preclinical trials: homozygous YACMJD84.2 (Q84/Q84)^31, 42^ and hemizygous CMVMJD135 (Q135)^43^ mice. We assessed two models to identify the common neurochemical abnormalities that may provide signatures of SCA3 pathology. To validate identified potential neurochemical biomarkers and determine which aspects of pathology they reflect, we evaluated cerebellar demyelination and axonopathy in the same mice, as well as brain tissue from patients with SCA3. Finally, we determined whether RNAi-mediated silencing of the SCA3 gene could reverse biomarkers of cerebellar demyelination and axonopathy in the Q84/Q84 model, since evidence of reversibility would suggest these noninvasive markers could reflect therapeutic efficacy in future trials.

## Materials and Methods

### Experimental design

The study was designed to assess MRS neurochemical profiles of two transgenic models frequently used in preclinical trials of candidate therapeutics for SCA3 – homozygous YACMJD84.2 (Q84/Q84)^42^ and hemizygous CMVMJD135 (Q135)^43^ mice. All animal procedures were approved by the University of Michigan Committee on the Use and Care of Animals (Protocol PRO00003836) and by the University of Minnesota Institutional Animal Care and Use Committee (Protocol # 1207A17510). Groups of Q84/Q84 (N=14 (7 females, 7 males), 9-17 month-old) and their littermate non-transgenic (wt) mice (N=11 (7 females, 4 males), 11-18 month-old), and Q135 (N= 9 (3 females, 6 males), 8-16 month-old) and their wt littermates (N=8 (5 females, 3 males), 10-16 month-old) in the C57Bl6/J background (Supplementary Table 1) were scanned. Mice were housed in cages with corncob bedding with a maximum number of five animals (range 3-5) and maintained in a standard 12-hours light/dark cycle with food and water *ad libitum*. Mice were shipped from the University of Michigan to the University of Minnesota for MR scanning, approximately 2-4 weeks prior to scans. After MR scanning, mice were deeply anesthetized with sodium pentobarbital (100 mg/kg i.p.), sacrificed by transcardial perfusion with pH 7.4 phosphate buffered saline and mouse tissue samples were sent back to the University of Michigan for molecular analysis.

### Mouse genotyping and CAG repeat size assessment

Genotyping was performed by PCR using DNA isolated from tail biopsy at the time of weaning as previously described^42, 43^. The CAG repeat size in Q84/Q84 and Q135 mice (Supplementary Table 1), and SCA3 patients and control individuals (Supplementary Table 2) was determined at Laragen Inc. by gene fragment analysis using primers hATAXN3forFam (5’ACAGCAGCAAAAGCAGCAA) and hATAXN3rev (5’CCAAGTGCTCCTGAACTGGT).

### Magnetic resonance (MR) protocol

Procedures for anesthesia and MR scanning were identical to our prior studies^44, 45^. Mice were induced with 3-4% isoflurane and were maintained anesthetized with 1.5-2% isoflurane during scanning. Body temperature was maintained at 36-37ºC and respiration rate at ~70-100 breaths per minute. The scanning time for each animal was approximately 50 minutes. MR scans were performed using a quadrature surface RF coil and a 9.4 T/31 cm magnet (Magnex Scientific, Abingdon, UK) interfaced to an Agilent console (Agilent, Inc., Palo Alto, CA, USA). A cerebellar volume-of-interest (VOI, 5-7 µl, Fig. 1A, 2A) was selected based on coronal and sagittal multi-slice images obtained with a rapid acquisition with relaxation enhancement (RARE) sequence^46^. All first- and second-order shims were adjusted using FASTMAP^47^. Localized ^1^H MR spectra were acquired with a short-echo localization by adiabatic selective refocusing (LASER) sequence (echo time TE = 15 ms, repetition time TR = 5 s, 256 averages)^48^, as described previously^44, 45^. Spectra were acquired and saved as single scans, which were individually frequency and phase corrected. Scans that showed evidence for motion were excluded and the remaining scans summed.

**Figure 1:**
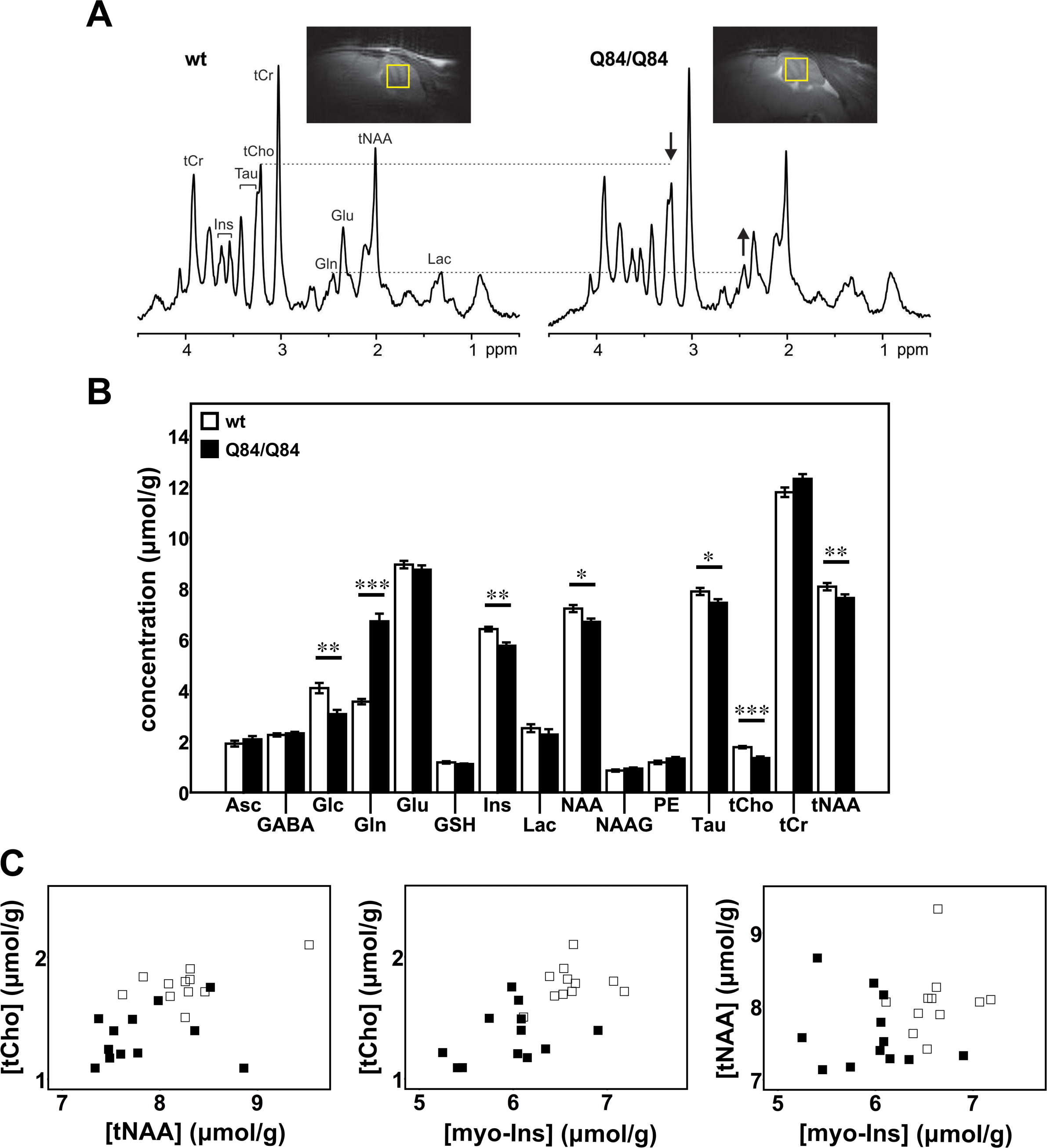
Cerebellar neurochemical levels are altered in homozygous Q84/Q84 mice. **A)** Representative localized proton MR spectra and midsagittal T_2_-weighted images of a Q84/Q84 and a wt littermate mouse. The most prominent neurochemical abnormalities, namely lower tCho and higher Gln in Q84/Q84 compared to wt mice, are denoted in the spectra by arrows. **B)** Cerebellar neurochemical profiles of Q84/Q84 mice (N=12, black bars) and wt littermates (N=11, white bars). Bars represent average neurochemical concentration ± SEM. Comparison between mouse genotypes was performed using Student’s t-test and statistical significance is indicated as: \**P*<0.05, \*\**P*<0.01, and \*\*\**P*<0.001. **C)** Separation of Q84/Q84 mice (black squares) from controls (white squares) by plotting two altered metabolites against each other.

### Metabolite quantification

The contributions of individual metabolites to the spectra were quantified using LCModel^49^ relative to unsuppressed water spectra acquired from the same VOI, as described before^44, 45^. Reliable concentrations were selected based on Cramér-Rao lower bounds (CRLB) criteria (quantification with CRLB ≤ 20% in the majority of the spectra). Alanine, aspartate, glycine and scyllo-inositol were excluded from final analysis based on these criteria. If the correlation between two metabolites was consistently high (correlation coefficient < - 0.5), their sum was reported^50^. Strong negative correlation was found between creatine and phosphocreatine and between glycerophosphocholine + phosphocholine, therefore total creatine (tCr) and total choline (tCho) were reported. Based on these reliability criteria,14 concentrations were evaluated: ascorbate (Asc), γ-aminobutyric acid (GABA), glucose (Glc), glutamine (Gln), glutamate (Glu), glutathione (GSH), *myo*-inositol (myo-Ins), lactate (Lac), *N*-acetylaspartate (NAA), *N*-acetylaspartylglutamate (NAAG), phosphoethanolamine (PE), taurine (Tau), tCr and tCho.

### Human Brain Samples

Deidentified frozen samples from the cerebellar cortex of autopsied brains from molecularly confirmed SCA3 individuals (N=3) and control individuals (N=4) (showing a CAG repeat size in *ATXN3* within the normal range) were obtained from the University of Michigan Brain Bank. Demographic information for these samples is provided in Supplementary Table 2.

### Western blot

Total proteins from mouse cerebella or human cerebellar cortex were obtained by lysis in RIPA buffer containing protease inhibitors (Complete, Roche Diagnostics), followed by sonication and centrifugation using an established protocol in the laboratory^31^. Supernatants (soluble protein fractions) were collected and total protein concentration was determined using the BCA method (Pierce). Total protein lysates from soluble fractions were resolved on 10% SDS-PAGE gels, and corresponding PVDF membranes were incubated overnight at 4 ºC with primary antibodies: rat anti-myelin basic protein (MBP) (1:1000; MCA409S, AbD Serotec), mouse anti-160 kD neurofilament medium (NFL) (1:1000; ab7794, Abcam), rabbit anti-α-tubulin (1:5000; 2144S, Cell Signaling), mouse anti-GAPDH (1:10000; MAB374, Millipore), and rabbit anti-MJD (1:20000). Bound primary antibodies were visualized by incubation with a peroxidase-conjugated anti-rat, anti-mouse or anti-rabbit secondary antibody (1:10000; Jackson Immuno Research Laboratories) followed by treatment with ECL-plus reagent (Western Lighting, PerkinElmer) and exposure to autoradiography films. Band intensity was quantified by densitometry using ImageJ.

### Statistical Analysis

Characteristics of transgenic and control groups were compared using Student’s t-test for age and chi-square test for sex. Levels of neurochemicals and proteins were compared between the transgenic and control groups using Student’s t-test as all distributions were normal and showed homogeneous variances. Linear regression analyses were performed to evaluate the relationship between neurochemical concentrations and protein levels using Pearson correlation. A *P* <0.05 was considered statistically significant in all analyses. Data were analyzed using IBM SPSS Statistics 22 software.

## Results

### Homozygous Q84/Q84 and hemizygous Q135 transgenic mice show shared neurochemical alterations in the cerebellum

Q84/Q84 and Q135 SCA3 transgenic mice show early onset of SCA3-like phenotypes starting around 6 weeks-of-age^31, 43^ and therefore are mouse models often used in preclinical trials of SCA3^31, 35, 43, 51–54^. We sought to investigate whether these two mouse models similarly altered neurochemical biomarkers that could serve as outcome measures in preclinical trials for evaluation of therapeutic efficacy.

Because no *in vivo* MRS studies were previously conducted in SCA3 mouse models, we obtained MR spectra at ultra-high field from the cerebellar vermis of symptomatic homozygous Q84/Q84, hemizygous Q135 and their littermate wt controls at an age when cell loss was previously shown in the cerebellum^43, 54^. Transgenic SCA3 mice were well-matched for age and sex to their wt control groups (*P*>0.05). Two Q84/Q84 mice, two Q135 mice and one Q135 wt control were found to have an abnormal high Gln profile that was previously described in the brains of mice derived from the C57BL/6 strain^55, 56^. The abnormal profile consisted of 2-3 fold higher Gln, 35-60% lower myo-Ins and 10-35% lower Tau relative to other wt or SCA3 mice in the same group. Because this profile results from portosystemic shunting in the liver^55^, which causes alterations in gene expression in many organs, these 5 mice were excluded from further analysis.

Neurochemical profiles of Q84/Q84 mice (N=12, 5 females and 7 males), with predominant CAG repeat size ranging from 69 to 74 triplets (Supplementary Table 1), showed significantly higher Gln and lower levels of total NAA (tNAA), NAA, Glc, myo-Ins, Tau, and tCho compared to wt littermates (N=11, 7 females and 4 males) (Figure 1A,B). To investigate if the Gln levels in Q84/Q84 mice that were almost two times higher than their wt controls were due to liver dysfunction, we analyzed serum liver profiles and liver pathology in a subgroup of mice.

This analysis revealed normal liver function and histology (data not shown) excluding liver dysfunction as the cause of the detected high levels of Gln in Q84/Q84 transgenic mice. Cerebellar neurometabolite profiles of Q135 mice (N=7, 3 females and 4 males), with CAG repeat length ranging from 125 and 129 trinucleotides (Supplementary Table 1), revealed significantly lower levels of tNAA, NAA, Glu, myo-Ins, GSH, and tCho compared to controls (N=7, 5 females and 2 males) (Figure 2A,B). Therefore, lower NAA and tNAA, tCho and myo-Ins represent the common neurochemical abnormalities in both models. In addition, groups of Q84/Q84 and Q135 mice were separated from wt controls with little or no overlap by plotting concentrations of these metabolites against each other (Figure 1C, 2C).

**Figure 2:**
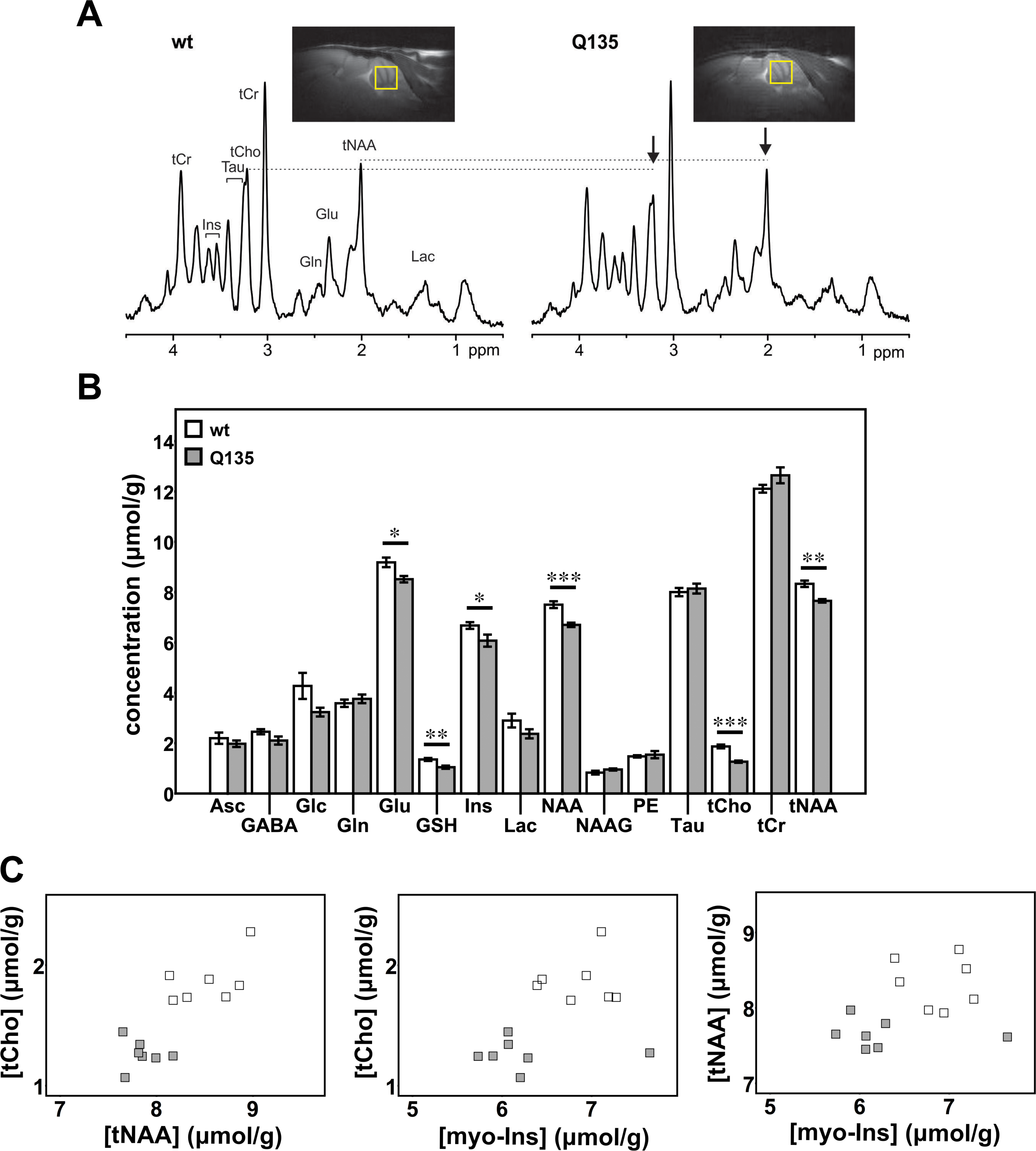
Cerebellar neurochemical levels are altered in hemizygous Q135 mice. **A)** Representative localized proton MR spectra and midsagittal T_2_-weighted images of a Q135 and a wt littermate mouse. The most prominent neurochemical abnormalities, namely lower tCho and tNAA in Q135 compared to wt mice, are denoted in the spectra by arrows. **B)** Cerebellar neurochemical profiles of Q135 mice (N=7, grey bars) and wt littermates (N=7, white bars). Bars represent average chemical concentration ± SEM. Comparison between mouse genotypes was performed using Student’s t-test and statistical significance is indicated as: \**P*<0.05, \*\**P*<0.01, and \*\*\**P*<0.001. **C)** Separation of Q135 mice (grey squares) from controls (white squares) by plotting two altered metabolites against each other.

### SCA3 transgenic mice show depletion of myelin basic protein (MBP) and neurofilament medium (NFL) in the cerebellum

Because the three neurochemicals that allowed clear separation of both SCA3 transgenic mouse models from their respective controls suggest disturbances in the metabolism of membrane and myelin phospholipids^57, 58^ (myo-Ins and Cho) and neuroaxonal pathology^59^ (NAA), and were all decreased in cerebella of SCA3 mice, we investigated whether these mice show cerebellar demyelination and axonal atrophy (Figure 3).

**Figure 3:**
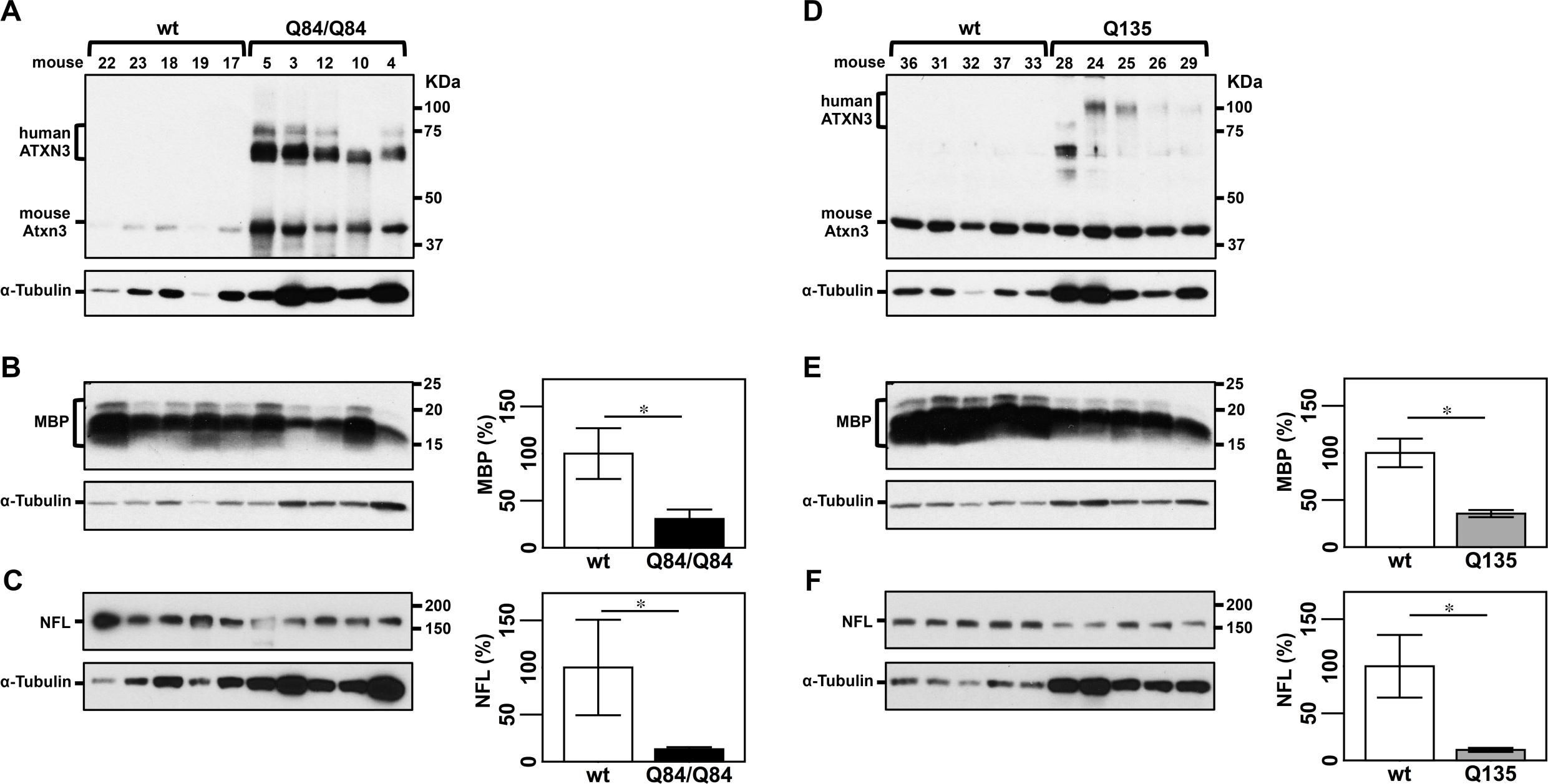
Cerebellar lysates from Q84/Q84 and Q135 mice show decreased levels of myelin basic protein (MBP) and neurofilament medium (NFL). **A,D)** Anti-ATXN3 immunoblot (anti-MJD antibody) detecting mutant human ATXN3 and endogenous mouse Atxn3 in cerebellar soluble protein extracts of the subset of Q84/Q84 (**A**), Q135 (**D**) and corresponding littermate wild type mice used to assess levels of MBP (**B,E**) and NFL medium (NFL) (**C,F**). Western blots show decreased levels of MBP and NFL in cerebella of Q84/Q84 (**B,C**) and Q135 (**E,F**) compared with controls. Graphs show quantification of protein bands by densitometry. Bars represent the average percentage of protein relative to respective wild type controls, normalized for α-Tubulin (± SEM). Statistical significance determined by Student’s t-test is indicated as \**P*<0.05.

Immunoblot analysis of cerebellar protein extracts from a subset (N=5 per group, 3 females and 2 males) of the mice used to obtain neurochemical profiles (Figure 3A,D) revealed that the levels of MBP were indeed reduced to 31% of wt levels in Q84/Q84 mice (Figure 3B) and to 35% of wt levels in Q135 mice (Figure 3E). NFL abundance was also reduced to 13% of wt levels in cerebella of Q84/Q84 mice (Figure 3C) and to 11% of wt levels in Q135 mice (Figure 3F).

### Levels of tCho and tNAA correlate with cerebellar abundance of MBP and NFL in SCA3 transgenic mice

We next sought to determine if levels of tCho and/or myo-Ins are indicators of the status of myelination and if tNAA levels reflect axon integrity in the cerebellum. Hence, we investigated correlations between the levels of these neurochemicals with abundance of MBP and NFL proteins in the two SCA3 mouse models (Supplementary Figure 1). Concentration of tCho correlated significantly with MBP levels in Q135 mice (*P*<0.001) (Supplementary Figure 1D) and showed a trend toward association in Q84/Q84 mice (*P*=0.165) (Supplementary Figure 1A). Levels of tNAA correlated significantly with NFL expression in Q135 mice (*P*=0.004) (Supplementary Figure 1F). No correlation was found between tNAA and NFL concentrations in Q84/Q84 mice (Supplementary Figure 1C), or between myo-Ins and MBP levels in either mouse model (Supplementary Figure 1B,E).

### MBP and NFL are lower in the cerebellar cortex of patients with SCA3 than healthy controls

To investigate whether dysregulation of cerebellar MBP and NFL in the two SCA3 transgenic mouse models reflects the human SCA3 condition, we assessed levels of these proteins in samples from cerebellar cortex of patients with SCA3 and healthy controls (Supplementary Table 2). Levels of MBP were decreased to 9% of control levels and NFL to 19% of control levels in cerebellar cortex of patients with SCA3 (Figure 4A-C) indicating demyelination and axonopathy in this brain region.

**Figure 4:**
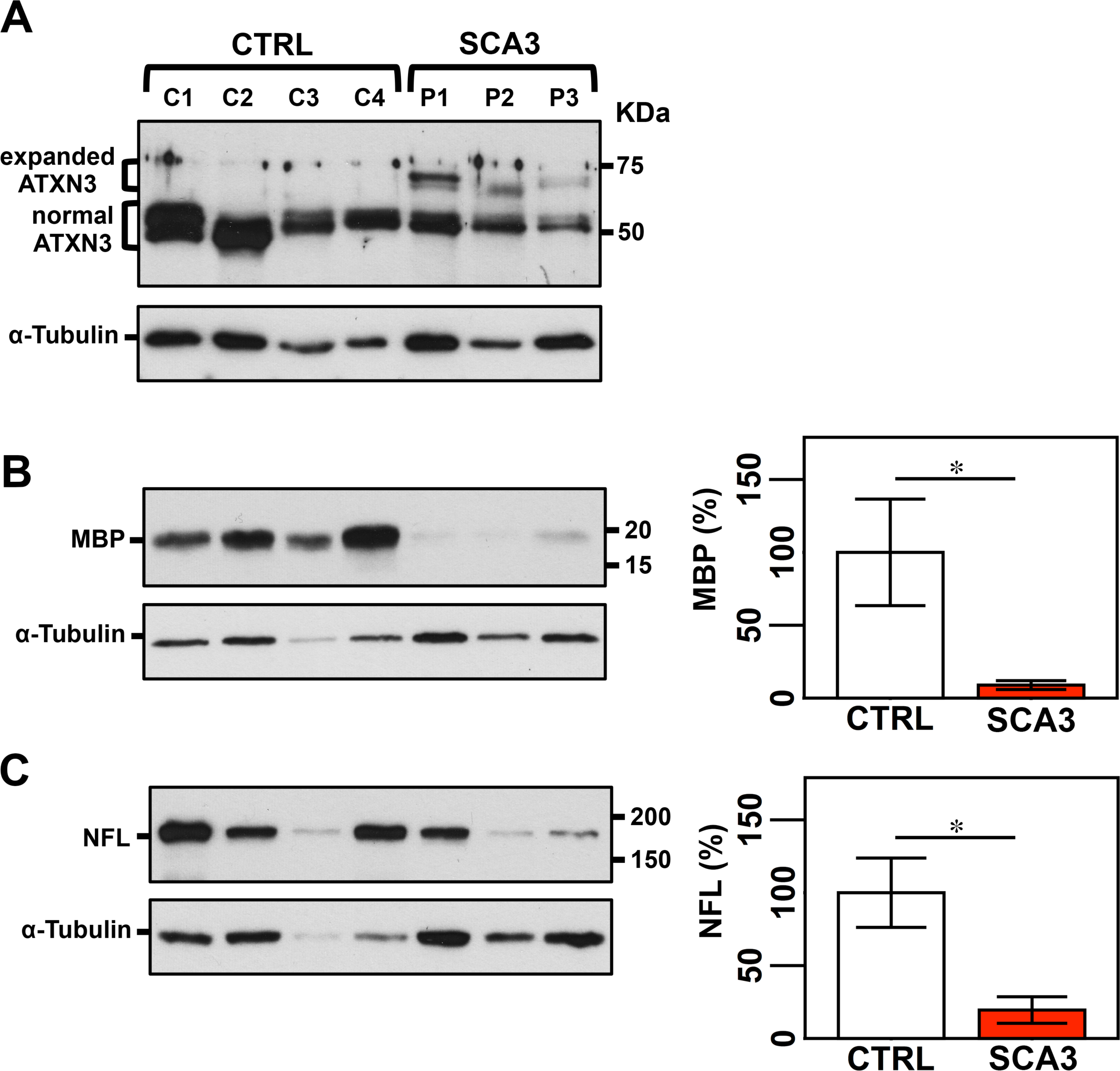
Protein lysates from cerebellar lobules of patients with SCA3 show decreased levels of MBP and NFL medium, indicative of demyelination and axonopathy/ neuronal loss. **A)** Anti-ATXN3 immunoblot (anti-MJD antibody) detecting expanded and normal ATXN3 in soluble protein extracts from cerebellar cortex of patients with SCA3 and control individuals. Detection of MBP (**B**) and NFL medium (**C**) by Western blotting shows lower levels of both proteins in patients compared with controls. Graphs show quantification of protein bands by densitometry. Bars represent the average percentage of protein relative to respective wild type controls, normalized for α-Tubulin (± SEM). Statistical significance determined by Student’s t-test is indicated as \**P*<0.05.

### *In vivo* long-term sustained reduction of mutant ATXN3 rescues cerebellar myelination and neurofilament biomarkers in homozygous Q84/Q84 mice

Because approaches to reduce levels of *ATXN3* gene products are currently a promising candidate therapy for SCA3, we sought to evaluate whether long-term sustained RNAi-mediated decrease of mutant human ATXN3 (hATXN3) abundance in Q84/Q84 cerebella would prevent reduction of MBP and NFL. Therefore, we used cerebellar protein extracts from end-stage Q84/Q84 mice that were injected at 6-to 8-weeks of age with an adeno-associated virus (AAV) encoding an artificial microRNA targeting mutant *ATXN3* (miRATXN3). We previously showed that miRATXN3 reduced hATXN3 levels to ~40% of vehicle-injected Q84/Q84 mice^31^. Cerebella of miRATXN3-treated Q84/Q84 mice in fact showed MBP and NFL levels of, respectively, 137% and 177% than their levels in vehicle-treated Q84/Q84 mice (Figure 5A,B). Whereas no association was detected between MBP and hATXN3 levels (*P*=0.218) (Figure 5C), NFL levels were significantly correlated with levels of mutant hATXN3, demonstrating an association between the level of gene silencing and the extent of rescue of neuroaxonal pathology in miRATXN3-treated Q84/Q84 mice (*P*=0.006) (Figure 5D).

**Figure 5:**
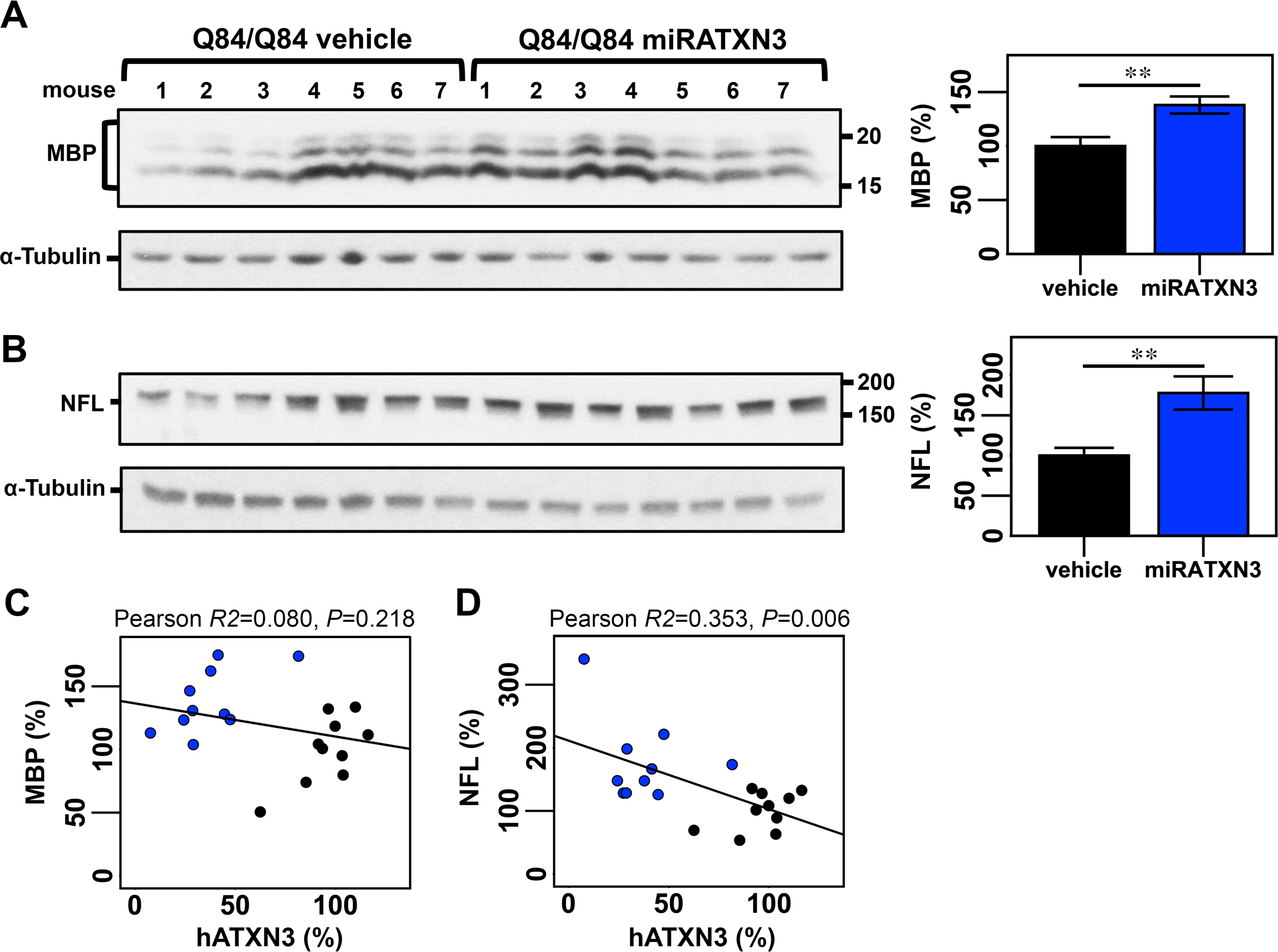
Cerebella from miRATXN3-treated Q84/Q84 show higher levels of MBP and NFL medium compared to vehicle-treated Q84 animals. Immunoblotting to detect MBP (**A**) and NFL (**B**) in cerebellar soluble protein extracts from end-stage Q84/Q84 mice injected with an adeno-associated virus delivering an artificial microRNA targeting ATXN3 transcripts (miRATXN3) or injected with vehicle. Graphs show quantification of protein bands by densitometry. Bars represent the average percentage of protein relative to levels in vehicle-injected Q84/Q84 mice, normalized for α-Tubulin (± SEM). Black and blue bars indicate, respectively, vehicle-injected and miRATXN3-injected Q84/Q84 mice. Statistical significance determined by Student’s t-test is indicated as \*\**P*<0.01. Plots showing Pearson correlations of levels of MBP with hATXN3 (**C**) and of NFL with hATXN3 (**D**) in vehicle (black circles) or miRATXN3 (blue circles) treated Q84/Q84 mice.

## Discussion

Future human clinical trials that seek to successfully translate preclinical animal studies would be facilitated by validated imaging biomarkers of cerebellar pathology. Here we identified shared neurochemical abnormalities in the homozygous Q84/Q84^31, 42^ and hemizygous Q135^43^ transgenic mice. Both of these models are well characterized models, reproduce several aspects of SCA3, and are extensively used in preclinical trials. We have further demonstrated associations of these noninvasive MR markers with protein markers of neuroaxonal pathology and demyelination, as well as the reversibility of these pathologies with mutant *ATXN3* gene silencing.

Previously cerebral and cerebellar metabolic alterations were only reported in one study that used *in vitro* ^1^H-MRS in brain tissue obtained from hemizygous Q84 mice^60^. The cerebellar neurometabolite spectra of homozygous Q84/Q84 mice showed similarities to the ones previously described in the *in vitro* study, namely higher Gln and lower NAA, tCho and myo-Ins in Q84 mice than wt controls^60^. Furthermore, the lower levels of tNAA observed in both models, lower levels of Glu in Q135 and higher levels of Gln in Q84/Q84 mice compared with controls, recapitulate prior observations in patients with SCA3^39, 41^. Decrease of tNAA in the cerebellum is a common biomarker of neuronal dysfunction or loss displayed by patients affected with several SCAs (SCA1, SCA2, SCA3/MJD, SCA6, and SCA7)^39, 41, 61, 62^, SCA1 mouse models^44, 45, 63^, and hemizygous Q84 mice^60^. The decreased Glu levels detected suggests dysfunction of glutamatergic granule cells, unipolar brush cells (UBCs), and/or parallel and climbing fibers in the cerebellar cortex^64^. While the number or function of these glutamatergic cells have not been investigated in cerebellar lobules of Q135 or Q84/Q84 mice, Purkinje cells (PCs) show decreased calbindin staining in Q135 mice^35^ and electrophysiological alterations in Q84 mice^54^ indicating that these cells are somehow dysfunctional. Because PCs receive excitatory input from parallel and/or climbing fibers, it is possible that Q135 PCs are impaired due to decreased input from these fibers into the molecular layer. End-stage Q84/Q84 cerebella did not show decreased levels of Glu, but displayed increased concentration of Gln, which could suggest abnormal Glu-Gln cycle between glutamate cells and Bergman glia in the molecular layer, and/or a higher number of glial cells that produce the bulk of Gln in the brain.

In addition to the above described neurochemical changes in SCA3 mice, each of the transgenic mouse groups showed a clear separation from their respective wild type controls in concentrations of tNAA, myo-Ins and tCho in the cerebellum. Because NAA is a validated marker of neuronal viability^59^ and a source of acetate for myelin lipid synthesis in oligodendrocytes^65^, myo-Ins is a structural sugar of membrane and myelin phospholipids^39^, changes in tCho levels reflect disturbances in phospholipid metabolism^65^, and all three metabolites were decreased in spectra of both SCA3 mouse models, we hypothesized that these mice would display cerebellar demyelination and axon atrophy. Demyelination, white matter and axon dysfunction and axonopathy have indeed been reported histopathologically in brain tissue obtained from patients with SCA3^17, 40, 41^. Furthermore, altered function of oligodendrocytes was recently reported in SCA3 models including the Q84^66^. Here, by showing that aged Q84/Q84 and Q135 cerebella and post-mortem cerebellar cortex from patients with SCA3 display reduced levels of MBP and NFL compared to their respective controls, we confirmed demyelination and axon loss in SCA3 cerebella. The magnitude of reduction we detected in NFL in SCA3 cerebella relative to controls was markedly higher than the reduction in cerebellar tNAA. Similarly, the magnitude of reduction in MBP in SCA3 was substantially higher than the % reduction in myo-Ins and tCho. This is likely because these neurochemicals have multiple roles, such as serving as metabolic intermediates and osmolytes, and therefore are less specific for neuroaxonal pathology than the protein markers. However, importantly, the associations between tCho with MBP levels and tNAA with NFL abundance, particularly in Q135 mice, suggest that tCho and tNAA levels reflect the status of myelination and axon integrity in the cerebellum of these mice. This evidence supports the inclusion of MRS evaluation of these neurochemicals during preclinical trials using Q84/Q84 and Q135 mice to evaluate the efficacy of therapeutic agents in rescuing SCA3 pathology.

While MBP levels were higher in miRATXN3-treated Q84/Q84 cerebella than wt controls, concentrations of MBP and hATXN3 did not correlate, possibly indicating that AAV2/1 miRATXN3 poorly transduced oligodendrocytes, as expected for this virus serotype^67^. Cerebellar axon loss seems to be, however, a consequence of expression of toxic expanded ATXN3 since reduction of NFL concentration in Q84/Q84 cerebella was suppressed by long-term decrease of mutant *ATXN3* levels in miRATXN3-treated Q84/Q84 mice. NFL levels significantly correlated with hATXN3 abundance in the cerebella of Q84/Q84 mice suggesting that AAV2/1 miRATXN3 effectively transduced and reduced hATXN3 in the majority of cerebellar neurons. Importantly, the reversal of the neuroaxonal pathology with RNAi-mediated reduction of ATXN3 levels supports the use of noninvasive imaging markers of this pathology in future pre-clinical and clinical trials that evaluate similar gene silencing strategies.

In summary, we have identified novel neurochemical and molecular similarities between end-stage homozygous Q84/Q84, aged hemizygous Q135 mice, and patients with SCA3, further validating the use of these mouse models in preclinical trials of SCA3. Like patients with SCA3, both transgenic models show neurochemical signatures indicative of neuronal loss, potential dysregulation of glutamatergic systems, and myelin and axon loss. We also provide molecular confirmation of cerebellar myelin and axon damage in SCA3 mice and patients, and establish significant associations of select neurochemical markers with invasive measures of myelin and axon damage. It will be important to perform future longitudinal MRS, molecular and pathological studies in parallel to evaluate the progression of neurometabolite dysregulation and associated molecular and pathological outcomes in the cerebellum and other SCA3-affected brain areas of transgenic mice. These novel neurochemical alterations in SCA3 transgenic mice reflect important aspects of pathology and should enable noninvasive monitoring of pathology reversal in future preclinical trials of therapeutic agents for SCA3.

## Acknowledgements

The authors thank the Michigan Brain Bank (5P30 AG053760 University of Michigan Alzheimer’s Disease Core Center) and its coordinator Mathew Perkins for providing brain samples of MJD patients and control individuals. This work was funded by a Becky Babcox Research Fund pilot research award, University of Michigan to M.C.C., NIH/NINDS R01NS03871 to H.L.P, and NIH/NINDS R21 NS111154 to H.S.M. and G.Ö. The Center for Magnetic Resonance Research is supported by the National Institute of Biomedical Imaging and Bioengineering (NIBIB) grant P41 EB027061, the Institutional Center Cores for Advanced Neuroimaging award P30 NS076408 and the W.M. Keck Foundation.

## Authors’ Roles

(1. Research project: A. Conception, B. Organization, C. Execution; 2. Statistical Analysis: A. Design, B. Execution, C. Review and Critique; 3. Manuscript Preparation: A. Writing of the first draft, B. Review and Critique)

M.C.C: 1) A, B, C; 2) A, B, C; 3) A, B

M.R.: 1) C

H.S.M.: 1) A, C; 3) B

N.S.A.: 1) C

S.F.: 1) C

V.G.S.: 1) A, B; 3) B

P.M.: 1) A, B; 3) B

H.L.P: 1) A; 3) B

G.O.: 1) A, B, C; 2) A, B, C; 3) A, B

