## Supplementary Material for "*In vivo* molecular signatures of cerebellar pathology in spinocerebellar ataxia type 3"

**Supplementary Table 1: Characteristics of transgenic mice used in this study.**

**Supplementary Table 2: Demographic information for postmortem human samples of cerebellar cortex used in this study.**

**Supplementary Figure 1: Neurochemical concentrations correlate with levels of MBP and NFL in Q135 mouse cerebella.** Plots showing Pearson correlations of levels of MBP with tCho (**A**,**D**), MBP with myo-Ins (**B**,**E**), and NFL with tNAA (**C**,**F**) in Q84/Q84 (black circles), Q135 (grey circles), and their respective wt littermate mice (white circles).
