## Supplementary figures and images for "*In vivo* molecular signatures of cerebellar pathology in spinocerebellar ataxia type 3"

### Supplementary Figure 1

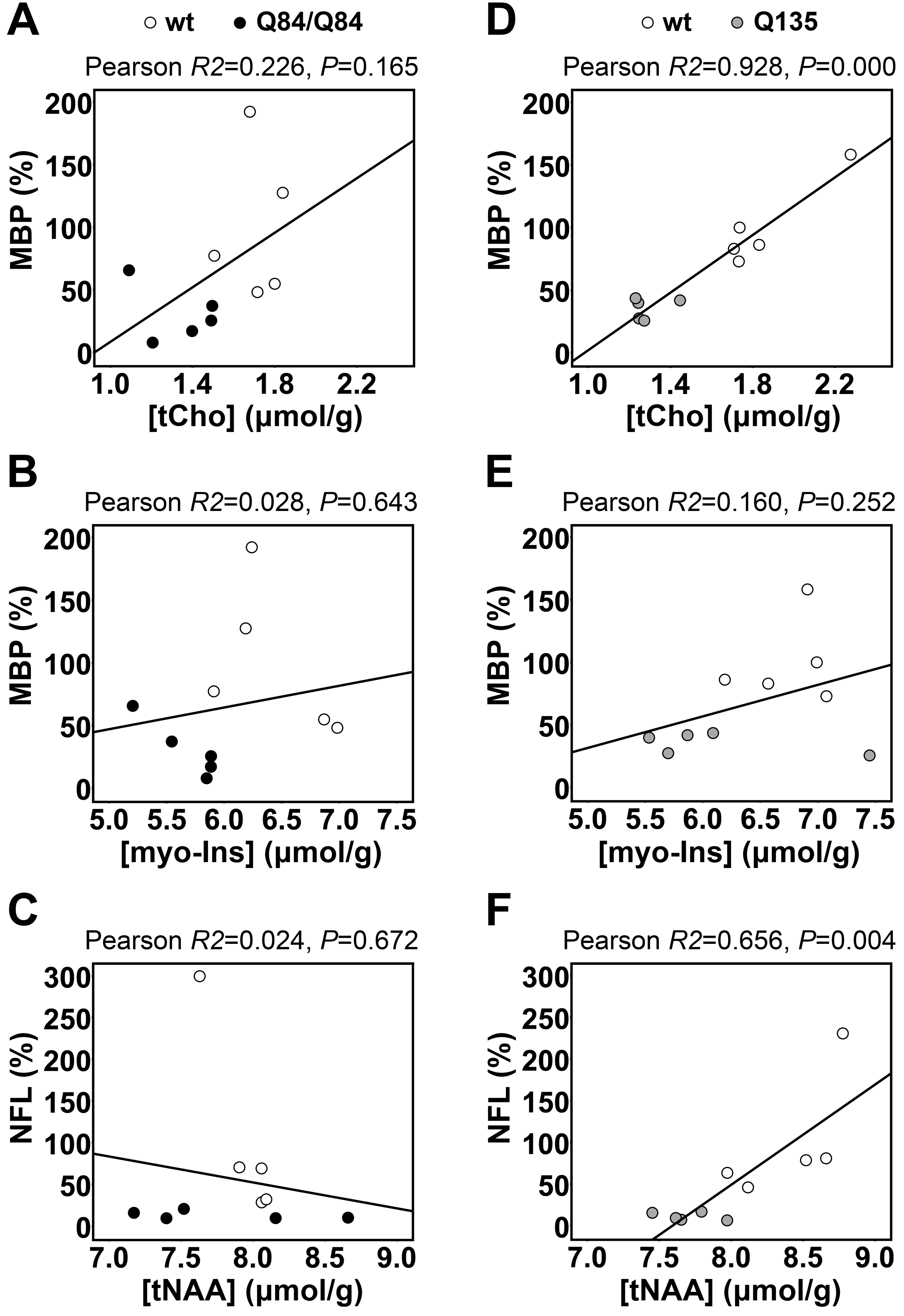
