## Supplementary Table 1 for "*In vivo* molecular signatures of cerebellar pathology in spinocerebellar ataxia type 3"

**Supplementary Table 1.** Characteristics of transgenic mice used in this study.

| Transgenic mouse line | Genotype | Sex | Mouse # | Age (months) | CAG repeat size(s) |
|---|---|---|---|---|---|
| YACMJD84.2 | Q84/Q84 | F | * | 12 | ND |
| | | | * | 10 | ND |
| | | | 1 | 12 | ND |
| | | | 2 | 11 | 70/**74**/80/87 |
| | | | 3 | 14 | **75** |
| | | | 4 | 17 | **70**/74/77/80 |
| | | | 5 | 15 | **70**/74/77/80 |
| | | M | 6 | 16 | ND |
| | | | 7 | 9 | **71**/77 |
| | | | 8 | 14 | **72**/78 |
| | | | 9 | 13 | **73**/77 |
| | | | 10 | 15 | **70**/73/77/81 |
| | | | 11 | 15 | **69**/75/76/82 |
| | | | 12 | 12 | 70/**74**/83 |
| | wt | F | 13 | 11 | NA |
| | | | 14 | 12 | |
| | | | 15 | 18 | |
| | | | 16 | 12 | |
| | | | 17 | 16 | |
| | | | 18 | 12 | |
| | | | 19 | 15 | |
| | | M | 20 | 13 | |
| | | | 21 | 13 | |
| | | | 22 | 15 | |
| | | | 23 | 14 | |
| CMVMJD135 | Q135 | F | 24 | 16 | **129** |
| | | | 25 | 16 | **127** |
| | | | 26 | 16 | **127** |
| | | M | * | 16 | **ND** |
| | | | * | 15 | **ND** |
| | | | 27 | 16 | **129** |
| | | | 28 | 16 | **125** |
| | | | 29 | 15 | **129** |
| | | | 30 | 8 | **129** |
| | wt | F | 31 | 16 | NA |
| | | | 32 | 16 | |
| | | | 33 | 16 | |
| | | | 34 | 16 | |
| | | | 35 | 10 | |
| | | M | * | 16 | |
| | | | 36 | 16 | |
| | | | 37 | 16 | |

Q84/Q84, homozygous YACMJD84.2; Q135, hemizygous CMVMJD135; F, Female; M, Male; NA, Not Applicable; ND, Not Determined; ∗, mouse eliminated from analysis due to detection of high levels of Glutamine; main transgene CAG repeat size is represented in bold.
