## Supplementary Table 2 for "*In vivo* molecular signatures of cerebellar pathology in spinocerebellar ataxia type 3"

**Supplementary Table 2.** Demographic information for postmortem human samples of cerebellar cortex used in this study.

| Health condition | Sex | Sample | Age at death | Cause of death | CAG repeat sizes |
|---|---|---|---|---|---|
| **Control individuals** | **F** | C2 | 48 | sudden cardiac arrest | 12/25 |
| | | C3 | 83 | polycythemia vera | 12/19 |
| | **M** | C1 | 59 | renal cell carcinoma | 18/21 |
| | | C4 | 61 | cardiac failure | 21/25 |
| **Patients with SCA3** | **F** | P1 | 59 | SCA3-related | 21/70 |
| | | P2 | 84 | | 21/66 |
| | | P3 | 48 | | 22/73 |

F, Female; M, Male
